# Map based cloning of *CT2* and the pilot functional exploration in abiotic stress

**DOI:** 10.1101/2024.03.17.585181

**Authors:** Chen Wang, Qi Zheng, Yi Xianggao, Zhanyong Guo, Lanjie Zheng, Jianping Yang, Jihua Tang, Weihuan Jin, Xu Zheng, Yong Shi

## Abstract

Heterotrimeric G-proteins are multifunctional modulators that participate in a wide range of growth and developmental processes in eukaryotic species, from yeast to plants and animals. Component detection and the study of G protein signaling in most plants, including maize, are in the initial stages. In this study, we characterized a maize mutant, *ct2*, that showed a compact architecture and reproductive organ-related phenotypic variation. The target gene *CT2* was cloned using bulked segregant analysis and map-based cloning. Gene structure prediction and phylogenetic analysis indicated that *CT2* is a canonical Gα protein belonging to the monocotyledonous group. Promoter analysis of *CT2* and RNA sequencing revealed *cis*-acting regulatory elements and differentially expressed genes involved in JA signaling and stress tolerance. The transcription of *CT2* was repressed by NaCl and PEG treatments, and *ct2* mutation in the *ct2* line compromised stress tolerance in maize. On the basis of our results, we proposed a schema diagram of *CT2*-regulated biological process and their feedback on *CT2* transcription. This research provides clues for further studies of *CT2* function in hormone signaling and stress tolerance, which is beneficial for maize breeding through the screening and application of beneficial alleles.

## Introduction

Plant growth and development are affected by heredity and the external environment. The basis of crop breeding is the manipulation of genetic material to improve target traits. Therefore, mining excellent gene resources and studying their mechanisms of action to improve crop genetics is necessary. Heterotrimeric GTP-binding protein (G protein) complexes, consisting of Gα, Gβ, and Gγ subunits, are multifunctional signal transducers involved in the growth, development, and transmission of environmental stimuli in all eukaryotes[1]. The mechanisms of action of G proteins were initially established in yeast and animals[2–4]. In metazoons, the signal transduction pathway begins with the activation of seven-transmembrane (7TM) G protein-coupled receptors (GPCRs) on the cell membrane by extracellular stimulation from ligand binding, such as hormones, neurotransmitter peptides, light, and sensory stimuli[5–7]. In the inactivated state, GDP binding Gα (Gα-GDP) interacts with heterodimer Gβγ to form Gαβγ complex. The binding of extracellular ligands activates GPCRs, which function as guanine nucleotide exchange factors to induce the exchange of Gα bound GDP for GTP(Gα-GTP). This leads to the dissociation of Gα-GTP and Gβγ. Gα-GTP and Gβγ are capable of relaying signals to downstream effectors through physical interaction[7–12]. The intrinsic GTPase activity of Gα hydrolyzes the Gα-bonded GTP to GDP, which could be accelerated by GTPase-activating proteins (GAPs); the resulting Gα-GDP then reassociates with Gβγ to reconstitute the inactive Gαβγ complex[10, 11, 13].

Subsequently, the molecular mechanism of G protein signaling has been elucidated in plants[2–4], benefitting from the completion of the sequencing of *Arabidopsis thaliana* and the identification of G protein mutants in *Arabidopsis,* rice, and maize[14–18]. The core components and signal transduction of G proteins are evolutionarily conserved in metazoas and plants[19, 20]. Plant G proteins participate in many growth and developmental processes, including cell division, plant architecture shaping, biological and abiotic stress tolerance, hormone signaling, sugar and light signal responses, nutrient use efficiency, and yield formation[20–22]. The signal transduction pathway of G proteins is extensive and agronomically significant; thus, the number of scientists in the plant field participating in the study of G proteins is increasing. Botella[23] asked a question—“Can heterotrimeric G proteins help feed the world?”—regarding the involvement of two Gγ subunits, grain size 3 (GS3), and dense and erect panicle 1 (DEP1), in the yield formation in rice. Notwithstanding conservation, the plant G protein system shows obvious divergence from that of animals[24]. For example, plant genomes encode significantly scarce subunits of each G protein, which does not affect its implication in many plant phenotypes[20]. The function of GPCRs is mainly exerted by signal receptor-like kinases with single transmembrane domains in plants, not by 7TM-GPCRs[25–29]. GPCR-like proteins in plants lack classical GEF activity but possess self-activating activity, and the G protein signaling is controlled through the level of protein deactivation [30–33]. Some phenotypes are regulated by canonical Gα subunits independently of nucleotide binding or heterotrimer dissociation[24, 34, 35]. Another discrepancy is the appearance of plant-specific extra-large G proteins (XLGs), which are also capable of binding to Gβγ heterodimers similar to canonical Gα subunits and participate in the regulation of growth and development in plants, such as vegetative growth, panicle and seed development, and tolerance to abiotic and biotic stresses[10].

Although the study of G protein signaling mechanisms in plants has substantially progressed, and the achievements have been reviewed in several papers[1, 10, 19, 36–40], further research is necessary. Maize, a major crop worldwide, has been used as a model for studying the evolution of plant nuclear genomes[41] and for basic and applied research in plant biology[42]. However, research on G proteins in maize is in the nascent stage. The maize genome encodes one canonical Gα (designated <u>C</u>ompact <u>P</u>lant 2; for clarity, herein, the name is CT2)[43], three XLGs (ZmXLG1, ZmXLG3a, and ZmXLG3b)[15], one Gβ and six Gγ proteins (including one TypeA, one TypeB, and four of the TypeC Gγ protein). Nevertheless, only the CT2[16, 43, 44], a Gγ proteins-ZmGS3[18, 45], and to some extent, 3 XLGs[15] were characterized. The functional research on CT2 has mainly focused on the effects on plant architecture, meristem development, and yield-related traits[16, 43, 44], excluding the other major functions of Gα in metazoans and plants, such as biological and abiotic stress resistance, hormone responses, cell division, and flower and panicle development[46–48]. Gene function mutants are useful for studying gene functions.

In this study, we report a maize <u>d</u>evelopmental <u>m</u>utant, *Zmdm* (designated *ct2* later), which displayed multiple phenotypes, including dwarf plant architecture; aberrant development of ears and tassels; and, most notably, yield-related traits. Using BSA followed by fine mapping with polymorphic <u>S</u>imple <u>S</u>equence <u>R</u>epeats (SSR) markers, the candidate gene was mapped to a 43.584 kb interval. By combining the RNA-sequencing data with subsequent genome sequencing and expression data, the target gene was identified as CT2. We conducted promoter analysis, gene structure analysis, transcriptome analysis, and an involvement study of *CT2* in signaling of phytohormones, salt or drought stresses, combining the diversity of plant architecture and yield-related traits between WT and *ct2* mutant, we proposed a schema diagram of the *CT2* regulated biological processes and their feedback to *CT2* transcription. This research provides clues for further studies of *CT2* function, which is beneficial for maize breeding, through the screening and application of beneficial alleles.

## Materials and Methods

### Plant materials and construction of the mapping population

*ct2*, a spontaneous mutant derived from the improved maize inbred line HCL645 (WT), was used as the female parent to pollinate L119 (a maize inbred line derived from the Huangzao4 heterosis group from China). The resulting F_1_ plants were self-pollinated or backcrossed as female parents with *ct2* as the recurrent parent to develop the F_2_ and BC_1_F_1_ populations, respectively. The resulting F_2_ population was used for inheritance analysis, and the BC_1_F_1_ group was adapted for BSA-seq and subsequent fine mapping of the target gene. The WT, *ct2*, F_2_, and BC_1_F_1_ populations were planted in a field at Yuanyang (35.07°N, 113.94°E), Henan Province, in the summer of 2022.

### Phenotyping of the *ct2* mutant

Maize plant height was measured from the ground to the top of the tassel. Stalk diameter was evaluated based on the diameter of the stem internode where the ear was born. The number of leaves per plant was calculated as the average total number of leaves per plant. Leaf length was defined as the average length from the collar to the tip. Leaf width was defined as the width of the widest part of the leaf blade. The tassel length indicated the distance from the uppermost internode to the top of the tassel. Rachis length was defined as the distance from the base of the uppermost tassel branch to the top of the rachis. Spikelet density was defined as the quantity of spikelets in the 2 cm region of the rachis 2 cm down from the top. At least 10 plants were investigated for each trait.

For ear-related traits, kernel row number, kernel thickness, kernel width, and kernel length were evaluated. Each trait was measured in three biological repeats with at least 10 ears or 15 kernels evaluated for each repeat. The average weight of three repeats (100 kernels per repeat) was used to evaluate the hundred-kernel weight.

### Whole genome sequencing and BSA-seq data analysis

Genomic DNA (gDNA) was isolated from maize leaves by using the CTAB method. A BC_1_F_1_ segregation population of 1577 individuals was used for BSA-seq. The WT and *ct2* plants were distinguished mainly by the phenotypes of dwarf and fasciated ear top in *ct2.* Two DNA pools (each with two independent replicates), constructed by bulking equal amounts of gDNA from 150 samples of either WT individuals (WT pool) or *ct2* mutant individuals (*ct2* pool), were sequenced on the Illumina Novaseq6000 platform at a sequencing depth of 50x at Beijing Berry Genomics. Raw data were filtered and analyzed according to the method of Zhang *et al*.[49]. In brief, to obtain clean reads, low-quality reads were removed and adaptors at both ends of the reads were trimmed using Fastp, as Chen *et al*.[50] with default parameters. The resulting clean reads were aligned to the reference genome of the maize inbred line B73 (B73_RefGen_v4) by using BWA software with SAM tools (Li and Durbin 2009). Polymorphic SNPs and InDels were detected using GATK software based on the comparison results between the sequences produced by sequencing and the reference sequences and annotated using ANNOVAR software. The candidate regions responsible for the mutant phenotype were estimated according to the ΔSNP index of each site, which indicated the difference between the ratio of SNP harboring reads to the entire number of reads of the same site (SNP-indices) of WT and mutant bulks.

### Fine mapping of candidate gene

The same BC_1_F_1_ segregation population used for the BSA-seq analysis was used for fine mapping of the mutation locus in *ct2.* SSR markers distributed between 26,335 and 75,373,051 bp on chromosome 1 were obtained either from the Maize Genetics and Genomics Database (maizeGDB: https://www.maizegdb.org) or manually searched using SSR hunter [51]. SSR markers were tested, 16 of which (Table S2) showed polymorphisms between *ct2* and L119 and were selected to identify recombined individuals.

### Real-time quantitative RT□PCR analyses

Total RNA was isolated from the leaves of maize seedlings by using TransZol reagent (ET121-01, TransGen Biotech, Beijing, China), following the manufacturers instructions. Removal of genomic DNA and reverse transcription of the mRNA were conducted using the *TransScript*^®^ One-Step gDNA Removal and cDNA Synthesis SuperMix kit (AT311-02, TransGen Biotech, Beijing, China). The resulting cDNA was used for qRT□PCR analyses using the *TransStart*^®^ Green qPCR SuperMix kit (AQ101-01, TransGen Biotech, Beijing, China) on a LightCycler® 480 System (Roche, Basel, Switzerland). *Ubiquitin 5* was selected as an internal control. Sequences of qRT-PCR primers for the target genes and internal control are provided in Table S2. Each qRT-PCR was performed in triplicate, and the average value of the relative expression (*Target gene*/ *Ubiquitin 5*) was used to represent the transcription levels of the corresponding genes. The transcriptional level of each gene was normalized to that of the WT background.

### PCR amplification of *CT2* and its promoter sequence

To amplify *CT2* and its promoter sequence, we designed the corresponding primers for the target sequences according to the reference sequence B73 (RefGen_v4) (Table S2). cDNA or genomic DNA samples isolated from leaves of WT or *ct2* line were used as templates and amplified by *TransTaq*^®^ DNA Polymerase High Fidelity (HiFi) (AQ131-11, TransGen Biotech, Beijing, China), followed by sequencing.

### RNA-seq and data analysis

Total RNA was extracted from leaves of 15-day-old seedlings of WT or *ct2*, and the method described for real-time quantitative RT-PCR analyses in this study was used. Each sample was prepared by pooling equal amounts of leaves from 10 plants of each genotype. mRNA was purified from total RNA by using polyT, followed by the construction of sequencing libraries. Libraries were prepared in two biological replicates and sequenced on the Illumina platform. Raw reads in the FASTQ format were processed using in-house Perl scripts and mapped to the Silva database to remove rRNA and obtain clean reads. Clean reads were mapped to the maize genome of B73 (B73_RefGen_v4) by using HISAT2[52]. The number of reads mapped to each gene was determined using Featurecount[53]. Differentially expressed genes (DEGs), which were defined as |log2(FoldChange)| ≥1 and P value<0.05, were evaluated by DESeq2[54]. GO analysis of the DEGs sets was conducted using the clusterProfiler package (https://bioconductor.org/packages/release/bioc/html/clusterProfiler.html, version 4.8.1) in R software (https://cran.r-project.org/) with a cutoff P value < 0.05.

### Phylogenetic Analysis of plant Gα proteins

Amino acid sequences of Gα proteins in *Zea mays* (maize), *Arabidopsis thaliana*, *Oryza sativa* (rice), *Triticum aestivum* (wheat), *Glycine max* (soybean), and *Solanum lycopersicum* (tomato) were aligned using ClustalW. A phylogenetic tree was constructed using MEGA X software (https://www. megasoftware.net/). The neighbor-joining method, based on 1,000 bootstrap repetitions, was used for construction.

### Preparation and microscopy of paraffin sections of maize seedling stems

Stem internodes (the fourth internode aboveground) of 4-week-old maize seedlings were sampled and cut into 2-3 mm thick segments in the transverse and longitudinal directions (near the central axis). The resulting segments were immediately placed in FAA fixative (G1101, Servicebio Technology Co., Ltd., Wuhan, China) for more than 24 h, stored, and transported at room temperature. Paraffin sections were prepared, and microscopy was performed by Servicebio Technology Co., Ltd., Wuhan, China. In brief, the method was as follows:1) Tissue fixation: tissues were removed from the fixative, smoothed using a scalpel in a fume hood, and then placed in the embedding frame with the corresponding labels. 2) Dehydration and wax leaching: the dehydration box (with the embedding frame) was placed into the dehydrator and processed: dehydrated with 75%, 85%, 90%, 95% alcohol for 4, 2, 2, and 1 h, respectively → anhydrous ethanol I and II, each for 30 min → alcohol benzene for 5∼10 min→ xylene I and II, each for 5-10 min → 65□ melting paraffin I, II and III, each for 1 h. 3) Embedding: the melted wax was placed into the embedding frame, tissues from the dewatering box were removed and placed into the embedding frame before the wax solidified according to the requirements of the embedding surface, and the corresponding labels were fixed. After Cooling at −20□ frozen platform, the wax block was removed from the embedding frame and repaired. 4) Section: The trimmed wax block was sliced into 4 μm slices. Tissue slices were flattened and picked up by the glass slides and baked in the oven at 60 □. After the water dried and the wax melted, the slides were removed and stored at room temperature. 5) Dewaxing and hydration: The paraffin sections were immersed in sequence in Environmentally Friendly Dewaxing Transparent Liquid (G1128, Servicebio) for 20 min, followed by → anhydrous ethanol for 5 min→ repeat former step → 75% ethyl alcohol for 5 min, rinsed with tap water, treated with Toluidine Blue for 2-5 min, rinsed with tap water, and dried in an oven. 6) Transparency and sealing: The sections were stained with xylene for 10 min and sealed with neutral gum, followed by microscopic inspection (upright optical microscope, NIKON ECLIPSE E100), image acquisition (imaging system, NIKON DS-U3), and analysis. Cell size in the transverse or longitudinal direction was evaluated using an average of more than 100 cells with three biological repeats.

### Measurement of Jasmonic acid levels

Maize kernels were sown in soil and grown in a growth chamber (14 h/10 h of light/dark, 30°C/25°C of day/night, 40%/60% relative humidity of day/night, 1,000 lx of light intensity), and the seedlings were grown to the 3-leaf stage for the follow-up experiments.

For determination of phytohormone content, the method was as follows:1) Metabolite extraction: The extraction of JA and asmonoyl-isoleucinewas (JA-Ile) followed the method descripted by Pan et al [55]. 2) Quantitative analysis of JA and JA-Ile. The contents of plant hormones in the samples are determined by Ultra High Performance Liquid Chromatography-Mass Spectrometry (HPLC-MS). The sample were separated by Agilent 1290 Infinity LC ultra-high performance liquid chromatography system. Samples were loaded in an automatic injector at 4□, the liquid chromatography column temperature was 45□. The mobile phase A was 0.05% formic acid aqueous solution, the mobile phase B was 0.05% formic acid acetonitrile solution. The flow rate was 400 μL/min, the sample size was 4μL. The relevant liquid phase gradient was as follows: 0-1min, the B phase changes linearly from 2% to 10%; 1—10min, the B phase changes linearly from 10% to 70%; From 10—11min, the B phase changed linearly from 70% to 95%. From 11—11.1min, the B phase changed linearly from 95% to 2%. From 11.1—13min, the B-phase was maintained at 2%. A QC sample was set up every certain number of experimental samples in the sample queue to detect and evaluate the stability and repeatability of the system. For correction of chromatographic retention time, standard mixtures of target substances was set up in the sample cohort. MS was performed in positive/negative ion mode on a 5500 QTRAP mass spectrometer (SCIEX). 5500 QTRAP ESI source positive ion conditions were as follows: source temperature: 550□; ion Source Gas1 (Gas1): 55; Ion Source Gas2 (Gas2): 50; Curtain. gas (CUR): 30; ionSapary Voltage Floating (ISVF): 4500 V. 5500 QTRAP ESI source negative ion conditions are as follows: source temperature: 550□; ion Source Gas1 (Gas1): 55; Ion Source Gas2 (Gas2): 50; Curtain gas (CUR): 30; ionSapary Voltage Floating (ISVF): −4500V. The ion pair was detected in MRM mode. 3) Analysis of data. The chromatographic peak area and retention time were extracted by Multiquant 3.0.2 software. The retention time was corrected by the standard of the target substance, and the contents of phytohormones were identified.

### Salt and drought stress tolerance experiments

The maize kernels of WT and *ct2* were germinated and hydroponically cultured with full-strength nutrient solution, as described by Geilfus *et al.*[56], and the culture condition was the same as that for plants for “Effect of phytohormones to *CT2* transcription.” Maize seedlings at the 3-leaf stage were treated with 200 sodium chloride (NaCl) or 5% polyethylene glycol (PEG) by adding NaCl or PEG6000 to the hydroponic solution at the corresponding concentrations. After 3 d of growing, the seedlings were phenotyped. Plant heights and fresh weights from at least 10 seedlings in each line were averaged and used to represent the plant heights and fresh weights of the corresponding lines.

### Measurement of MDA contents, SOD, and POD activity

Culture conditions of maize seedlings were the same as those for "seedling for salt and drought stress tolerance experiment" in this paper. The leaves of maize seedlings at the 3-leaf stage were sampled. MDA content and SOD and POD activities were estimated using the methods of Zhang *et al.* [57] (for the determination of or the determination of content) and Giannopolitis *et al*. [58] (for the determination of SOD and POD activity). At least 10 seedlings were measured, and the average value was calculated for each trait.

## Results

### Map-based cloning of the candidate gene resulting in the *ct2* mutant phenotype

The *Zmdm* mutant, a spontaneous mutated inbred line in HCL645 (WT) background, was screened as a <u>d</u>warf <u>m</u>utant among the breeding materials. The candidate gene underling the mutant phenotypes have been identified as maize Gα subunit – *CT2,* later. To avoid confusion, we redesignated the *Zmdm* mutant as *ct2*. *ct2* plants were dwarfed from the vegetative stage to the reproductive stage, with a plant height of only 57% of that of WT (Figure 1A, B, C, E). The leaves of *ct2* were significantly shorter than those of WT, but the leaf width remained unchanged (Figure 1D, F, G). To describe the genetic characteristics of *ct2*, we crossed *ct2* (as *a* female parent) with the inbred maize line L119, which belongs to the Huangzao4 heterosis group from China. The resulting F_1_ plants were self-pollinated or backcrossed as the female parent with *ct2* to obtain the F_2_ or BC_1_F_1_ populations. The F_1_ plants developed normally, and the F_2_ population showed approximately 3:1 segregation (157:48, χ^2^=0.1967, *p*=0.6) of WT versus mutant plants, indicating that the mutant phenotypes resulted from monogenic inheritance. This was verified in the BC_1_F_1_ population (1577 individuals), which showed a separation ratio of WT versus *ct2* close to 1:1 (775:802, χ^2^=0.4287, *p*=0.5). The same BC_1_F_1_ population was used to map candidate genes. The WT and *ct2* plants were distinguished mainly by the phenotypes of dwarf and fasciated ear top in *ct2.* We constructed two DNA pools (each with two independent replicates) by bulking equal amounts of gDNA from 150 samples of either WT or *ct2* mutant individuals and sequenced them for BSA. The results preliminarily mapped the candidate gene to a region between 26,335 and 75,373,051 bp on chromosome 1 in the maize genome (Table S1).

**Figure 1.**
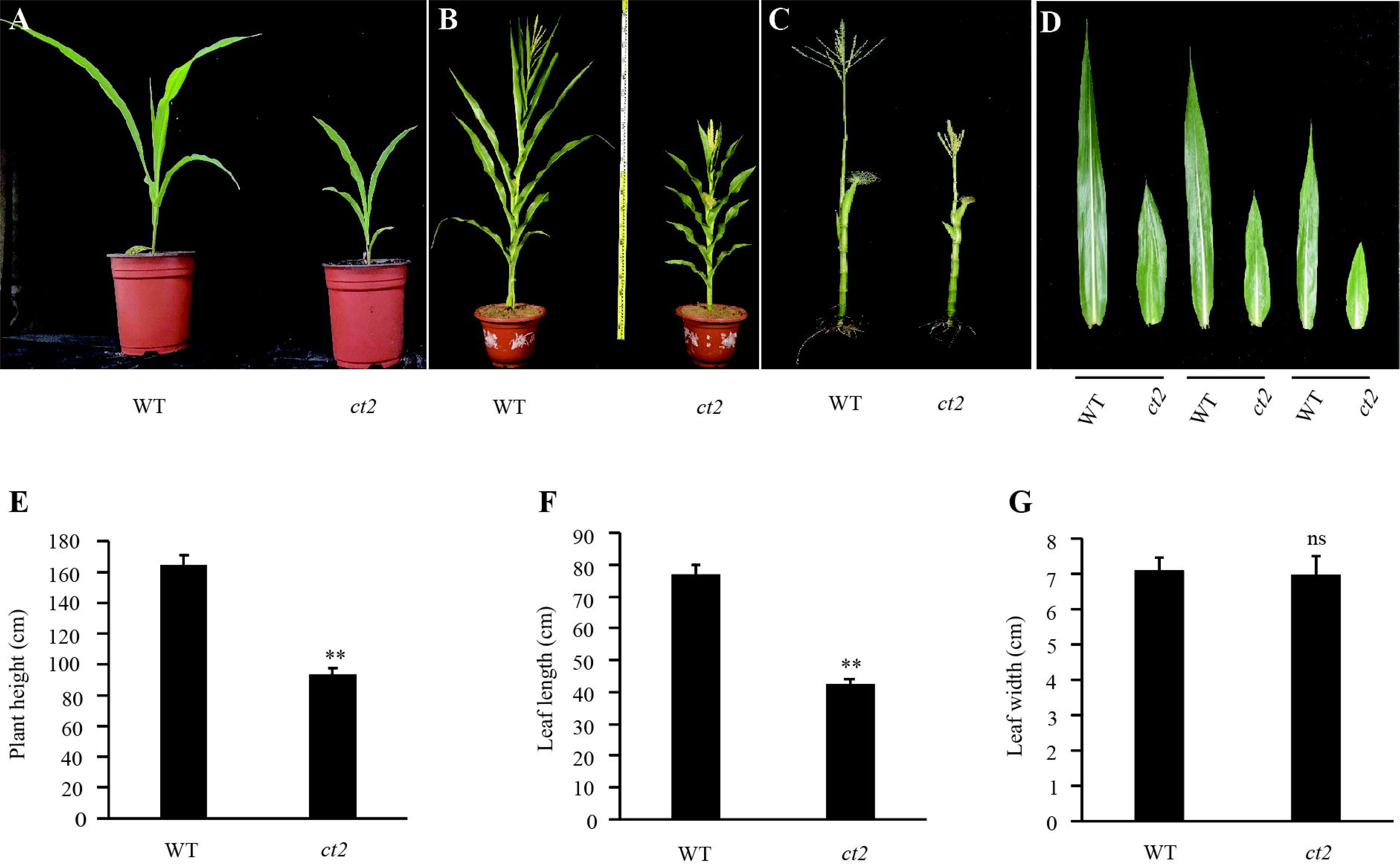
Plant phenotypes of *ct2* mutant. **A**, **B**, *ct2* mutants were dwarfed at both juvenile stage (3-leaf stage) **(A)** and mature stage **(B)**. **C**, Plant phenotype of WT and *ct2* mutant with leaves striped at mature stage. **D**, Phenotype of antepenultimate leaf, penultimate leaf and apical leaf in WT and *ct2* mutant. **E-G**, Quantification of plant height **(E)**, leaf length **(F)** and leaf width **(G)** of antepenultimate leaf in WT and *ct2* mutant. Error bars represent stand deviation errors. ns, * and ** indicate significant levels of not significant, P<0.05 or P<0.01, respectively, by single-tailed students *t* test.

Next, we fine-mapped the candidate gene to a 43.346 kb region between SSR makers 16,707,270 and 16,750,0616 by using common and custom SSR markers (Figure 2A). Depending on the annotation of the reference genome of maize inbreed line B73 (B73_RefGen_v4), five putative genes (Zm00001d027884, Zm00001d027885, Zm00001d027886, Zm00001d027887, Zm00001d027888) were concluded in this region. RNA-sequencing (RNA-seq) analysis indicated that only the transcription level of Zm00001d027886 (*CT2*) was significantly reduced (Table S3) in *ct2* compared with WT, verified by qRT-PCR analysis (Figure 2B). To determine the underlying mechanism, we amplified and sequenced the promoter regions of *ct2* and WT. Three transversions (T->C, A->G, C->A at −2000, −864, and-832, respectively) and a single nucleotide (cytosine at −1487) deletion before the translation initiation codon (ATG) of *CT2* were detected (Figure S1). However, that the decreased transcription of *CT2* resulted from these variations is unlikely because the promoter sequence of *ct2* (−2320->-1 from ATG) in the mutant was identical to that of B73, which represents normal plant height and reproductive organs.

**Figure 2.**
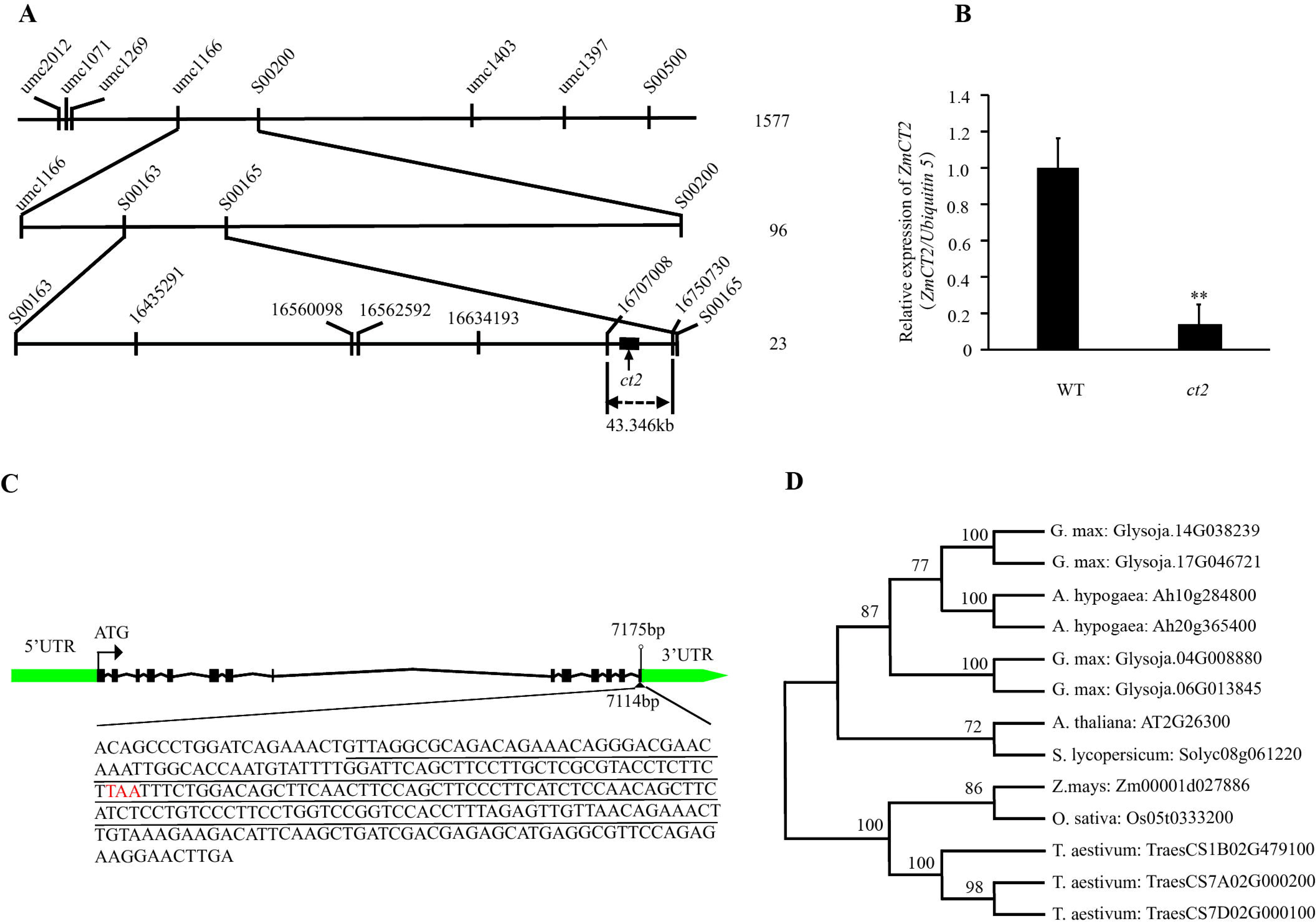
Fine mapping, transcriptional analysis, gene structure and phylogenetic analysis of *CT2* (or the putatively encoded amnio acid sequence) **A**, Sequential fine mapping of *CT2*. BC_1_F_1_ segregation population was employed to map the candidate gene. The candidate gene was preliminary mapped to a region between 26,335bp to 7,537,3051bp on chromosome 1. SSR makers used for fine mapping and numbers of plants in each mapping step (on the right) were indicated. **B**, Expression analysis of *CT2* in leaves of maize seedlings at 3-leaf stage by qRT-PCR. *Ubiqutin5* gene was used as the internal control. Relative transcriptional levels of *ZmCT1* were normalized to the expression level in WT. Error bars represent stand deviation errors. ** indicates significant level of P<0.01 by single-tailed student’s *t* test. **C**, Schematic representation of *CT2* structure. The 5’, 3’ untranslated regions (5’, 3’ UTR), exons and introns were indicated in green rectangle, green arrow, black rectangles and black broken lines, respectively. The translation initiation site (ATG), the site of the 185bp insertion (7114bp form ATG), sequence of 185bp insertion in *ct2* mutant (underlined), the site of the stop codon (TGA, 7175bp from ATG) and the nonsense mutation of the stop codon (TAA, highlighted in red) in the insertion sequence were also annotated. **D**, Phylogenetic tree of CT2 in *Zea mays* (maize), *Arabidopsis thaliana*, *Oryza sativa* (rice), *Triticum aestivum* (wheat), *Glycine max* (soybean), *Solanum lycopersicum* (tomato). The phylogenetic tree was constructed by MEGA X using the neighbor-joining method based on 1,000 bootstrap repetitions.

Next, we amplified the cDNA sequences of the five genes included in the candidate 43.584 kb region in *ct2* and WT backgrounds. The sequence comparison showed that only the CDS of *CT2* line encoded a mutated amino acid sequence in *ct2* with putatively amino acid sequences encoded by other four genes keeping unchanged. The mutation in *ct2* resulted from an 185 bp sequence insertion after the 1112 bp (after 7114 bp in genomic DNA) in wild-type *CT2*, which led to a 61 bp replacement by an 84 bp sequence at the 3’termial of *CT2* (Figure 2C, S2).

### Gene structure and promoter analysis and of *CT2*

We next performed the gene structure analysis of *ZmCT2* and the putatively encoded protein. The genomic structure of *CT2* is complex, with 14 exons and 13 introns distributed throughout a 7175 bp chromosome segment (Figure 2C). The structure of CT2 was predicted using InterPro software (http://www.ebi.ac.uk/interpro/). The results indicated that CT2 is a canonical Gα protein with a receptor binding site, GoLoco binding site, switch I and II region, βγ complex interaction site, GTP/Mg2+ binding site, and adenylyl cyclase interaction site (Figure S3). Although G proteins in plants share structural similarities with their animal counterparts, their signaling mechanisms vary substantially [22]. Functional variations exist even in the plant kingdom [18, 45, 59, 60]. Thus, a phylogenetic analysis of Gα in common monocotyledonous and dicotyledonous crops, including *Zea mays* (maize), *Arabidopsis thaliana*, *Oryza sativa* (rice), *Triticum aestivum* (wheat), *Glycine max* (soybean), *Solanum lycopersicum* (tomato), was conducted. Unlike in animals, all plants included here encode no more than 4 Gα proteins. This result corresponds well with those of Urano *et al.* [22, 33]: the number of G protein components and the RGS (G Signaling proteins) varied in different plant species and were detected or deleted together in the given genome. The evolutionary tree showed that the homology of Gα is higher within monocotyledonous or dicotyledonous plants than between them (Figure 2D), which may indicate a bifurcation before the divergence of monocotyledonous and dicotyledonous plants. The *CT2* was classified to the monocotyledonous group.

To understand and predict the physiological function of *CT2*, we submitted its promoter sequence (from −2320 to 0 bp before the translation initiation site ATG) to the Plant CARE website (http://bioinformatics.psb.ugent.be/webtools/plantcare/html/) to detect cis-acting regulatory elements and found that the promoter of *CT2* contains sites involved in defense and stress response, anaerobic induction, light response, abscisic acid (ABA), jasmonic acid methyl ester (MeJA) response, and meristem expression (Table S4). Thus, we conclude that *CT2* is a complex gene and encodes a typical Gα protein, which may be involved in the pathway of meristem development, phytohormone regulation, environmental signal perception, and stress response.

### *ZmCT2* is involved in development of reproductive organs in maize

CT2 has been shown to participate in the *CLAVATA (CLV)-WUSCHEL (WUS)* signaling pathway. Mutations in CT2 result in fasciated meristems and ears[43, 61]. *ct2* also showed developmental defects in both male and female organs. The tassels of ct2 were shorter and compact with thicker rachises and branches (Figure 3A, B, I, J). Spikelets on ct2 tassels were compact, and the spikelet density of rachis and branches increased to over 1.7 times that of WT (Figure 3C, K, L). The top of the young ear was slightly fasciated in *ct2*. However, fasciation was hardly detected in the mature ears (Figure 3D, E). The ears of *ct2* were shorter than those of WT, but for the former, the kernel row number was unaffected (Figure 4S4M). Kernel thickness, kernel width, and kernel length of *ct2* ears were 154%, 100%, and 70% those of WT (Figure 3F-H, M-O). The hundred-kernel weight of *ct2*, which was 92% that of WT, declined significantly (Figure 3Q). In summary, *ct2* exhibited compact plant architecture and reproductive organs, and we conclude that CT2 is involved in maize growth and development, including SAM development.

**Figure 3.**
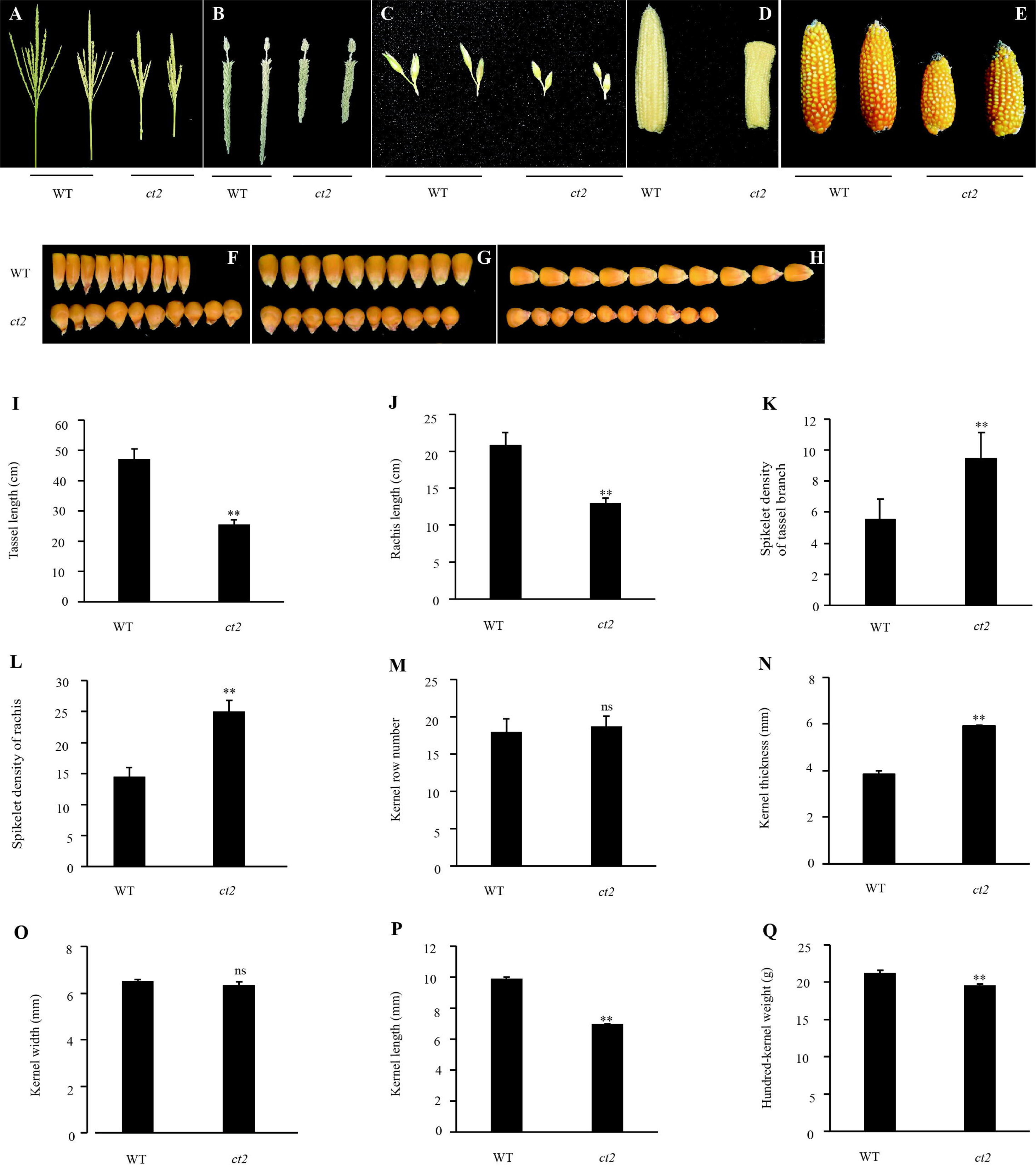
Phenotyping of reproductive organ related traits in *ct2* mutant. **A**, **B**, **C**, Tassels **(A)**, tassel rachis **(B)**, tassel spikelet **(C)** in WT and *ct2* mutant. **D**, Top of young ears of *ct2* are fasciated. **E**, Matured ear of WT and *ct2* mutant. **F-H**, Kernel thickness **(F)**, kernel width **(G)** and kernel length **(H)** of WT and *ct2*. **I-Q**, Quantification of tassel length **(I)**, rachis length **(J)**, spikelet density of tassel branchs **(K)**, spikelet density of rachis **(L)**, kernel row number **(M)**, kernel thickness **(N)**, kernel width **(O)**, kernel length **(P)**, hundred-kernel weight **(Q)** in WT and *ct2* mutant. Error bars represent stand deviation errors. ns, * and ** indicate significant levels of not significant, P<0.05 or P<0.01, respectively, by single-tailed student’s *t* test.

**Figure 4.**
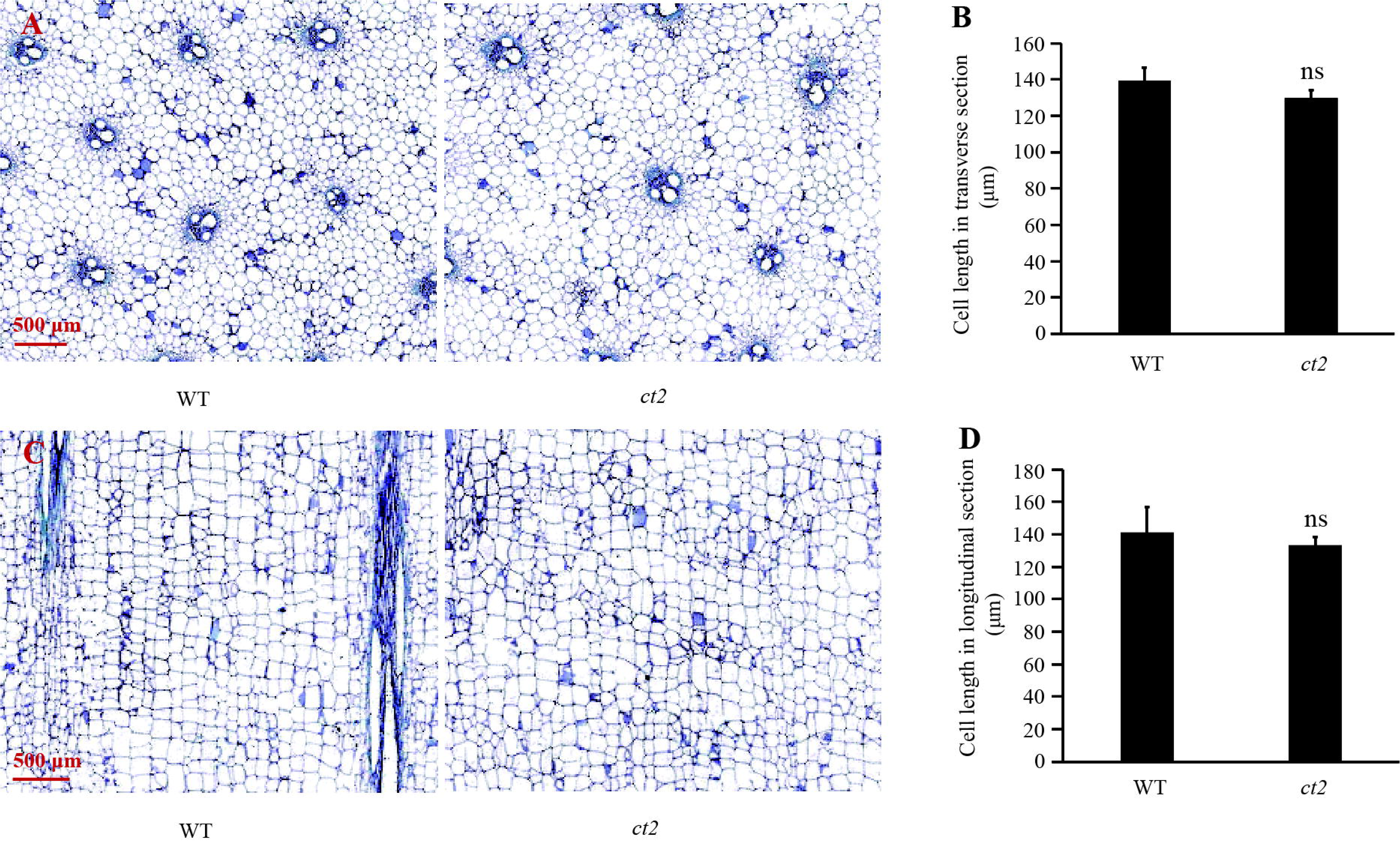
Sizes of stem cells in WT and *ct2* mutant seedlings. Paraffin section in transverse or longitudinal direction of stem internodes of 4-week maize seedlings in WT and *ct2* mutant were inspected through microscopy. **A**, **B**, Cells of stem internodes in transverse direction in WT (left panel) and *ct2* mutant (right panel) **(A)** and the cell lengths were quantified **(B)**. **C**, **D**, Cells stem internodes in longitudinal direction in WT (left panel) and *ct2* mutant (right panel) **(C)** and the cell lengths were quantified **(D)**. Error bars represent stand deviation errors. ns, * and ** indicate significant levels of not significant, P<0.05 or P<0.01, respectively, by single-tailed student’s *t* test. Different lowercase letters indicate significant differences (*P < 0.05) by one-way analysis of variance (ANOVA) and post hoc Duncan test.

### *CT2* mutation inhibits the elongation of stalk internode

The null mutation of *ct2* reduces plant height but increases the total leaves per plant [16]. The *ct2* mutant described in this study also exhibited serious dwarf phenotypes (Figure 1A, B, C, and E), and the stem diameter and number of leaves per plant were not affected (Figure S4). However, Urano *et al.* (2015) showed that the mutation of *CT2* increased the total number of leaves. The discrepant results may result from that Urano *et al.* recruited an *ct2* null mutant, the mutant descripted here could still encode a truncted *ct2*. To determine the mechanism underlying reduced plant height, we examined the cell size of maize seedling stem paraffin sections from WT and *ct2* backgrounds by microscopy. The results showed that cell size in the stalk internode of *ct2* was not affected in either the transverse or longitudinal direction compared with that in WT (Figure 4). Because the stem diameter and number of leaves per plant in *ct2* are equal to those in WT (Figure S4) and *CT2* modulates cell proliferation of shoots and roots in maize [16], we speculated that the reduced plant height in *ct2* resulted from the reduction in cell numbers, which may have resulted from the retarded cell proliferation in stalk internodes.

### RNA-seq and data analysis of *ct2*

To further understand the biological function of *CT2*, the leaves of a 15-day-old seedling of WT and *ct2* mutant were sampled, followed by mRNA sequencing (mRNA-seq). Gene transcription levels were quantified with the threshold value of |log_2_(FoldChange)| ≥1 and significance score P value < 0.05. Globally, the transcription levels of 1156 genes were altered, with 462 upregulated and 694 downregulated genes in *CT2* compared with WT (Table S3). Gene ontology (GO) analysis of the downregulated genes was performed, and the top 20 terms in each category are listed in Figure 5. The major functions in the biological processes of the 694 downregulated genes included JA signaling, response to wounding, response to stress, defense response, regulation of signal transduction, and regulation of cell communication, indicating that *CT2* may be involved in phytohormone (JA) signaling, stress, and signal transduction.

**Figure 5.**
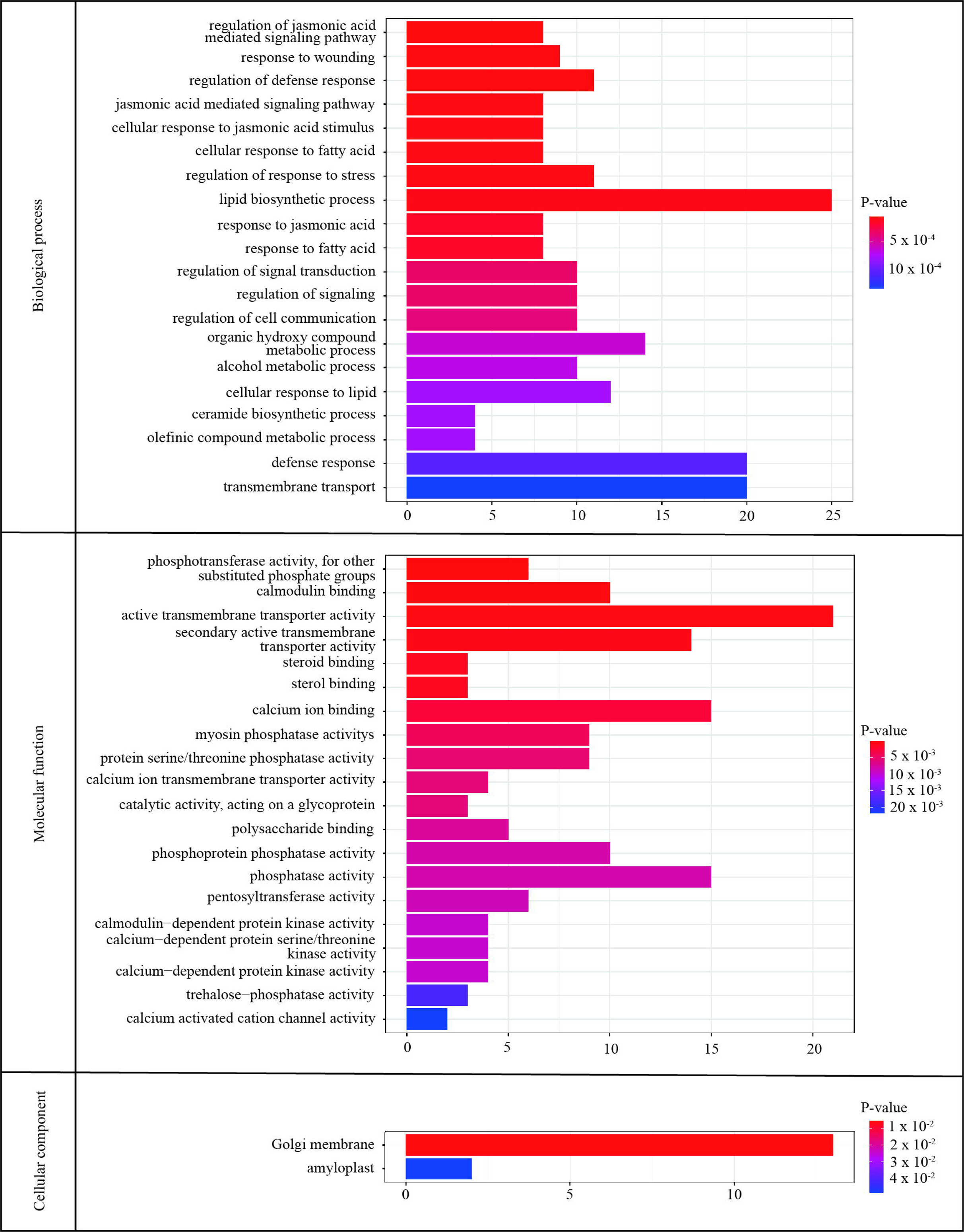
Gene ontology analysis of significantly downregulated genes in *ct2* mutant compared to WT. Total RNA was extracted from leaves of 15-day-old maize seedlings followed by deep sequencing and data analysis. The differentially expressed genes (|log2(FoldChange)| ≥1 and P-value <0.05) were submitted for gene ontology analysis.

### *CT2* positively regulates Jasmonoyl-isoleucine accumulation and involved in salt and drought tolerance in maize

G proteins interact with hormones to fine-tune plant biological processes[62] and are involved in plant biotic and abiotic stress responses[19]. In addition to the occurrence of JA (Jasmonic acid)-responsive cis-regulation in *CT2* promoter and DEGs in JA signaling and stress responses in GO analysis, we aimed to determine the interaction of *CT2* with phytohormones or stress tolerance. We compared the phytohormone levels of JA, Jasmonoyl-isoleucine (JA-Ile) in WT and *ct2* background. The results represented *ct2* mutation in *ct2* signficantly depressed the contents of JA-Ile (Figure 6 A, B).

**Figure 6.**
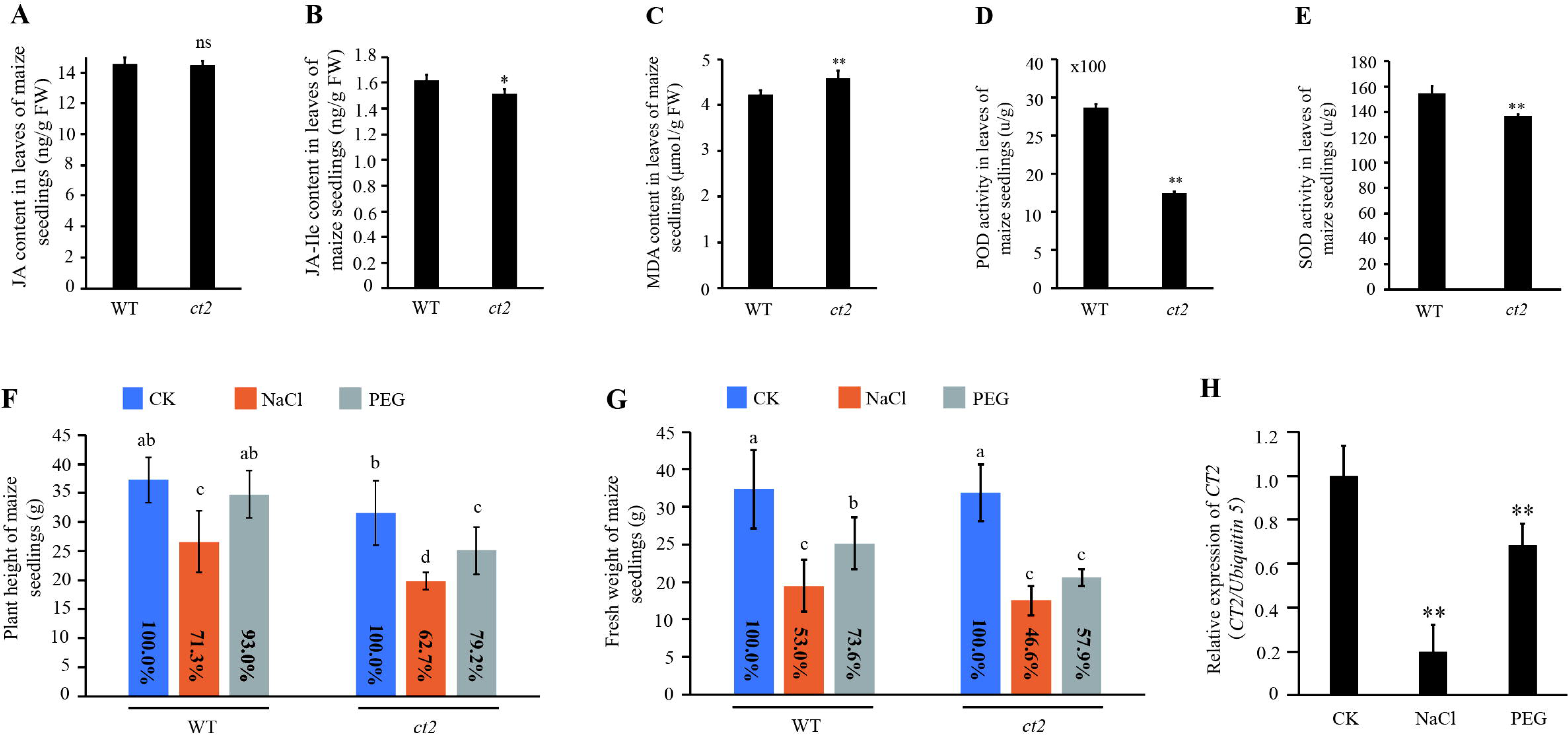
Involvement of *CT2* in phytohormone signaling and stress tolerance Seedlings at 3-leaf edge were used for experiments. For phytohormone treatment, seedlings were foliage sprayed with ABA, IAA, GA3, 6-BA, BR or MJA and the leaves were sampled 1 hour after phytohormone treatment for the following mRNA isolation and RT-PCR analysis. For salt and drought stress treatment, hydroponic cultured seedlings were treated with NaCl or PEG then were collected for further experiment after 3 days culturing. **A, B,** Jasmonic acid (JA) **(A)**, Jasmonoyl-isoleucine (JA-Ile) **(B)** contents in WT and *ct2* seedlings. **C, D, E**, MDA contents **(C)**, POD activity **(D)** and SOD activity **(E)** in WT and *ct2* seedlings. **F**, **G**, Effect of plant height **(F)** and fresh weight **(G)** of WT and *CT2* seedlings under salt and drought treatment. Percentage values relative to wild-type controls are indicated. **H,** Expression analysis of *CT2* in WT seedlings under salt and drought treatment by qRT-PCR. Error bars represent stand deviation errors. ns, * and ** indicate significant levels of not significant, P<0.05 or P<0.01, respectively, by single-tailed student’s *t* test. Different lowercase letters indicating significant differences (*P < 0.05) by one-way analysis of variance (ANOVA) and post hoc Duncan test.

Environmental stress induces the production of reactive oxygen species (ROS), and plants have evolved strategies to scavenge these oxidative compounds, including the expression of ROS-scavenging enzymes[63]. The examination of malondialdehyde (MDA) content, superoxide dismutase (SOD), and peroxidase (POD) activity in WT and *ct2* showed that mutation in *ct2* resulted in the upregulation of MDA content and downregulation of SOD and POD activity (Figure 6C-E), which indicated that *CT2* mutation may decrease stress tolerance in *ct2*. To test this hypothesis, we treated hydroponically cultured WT and *ct2* seedlings with NaCl (200mM) or PEG6000 (5%) to mimic salt and drought stress. The NaCl and PEG treatments repressed the growth (plant height and fresh weight) of WT seedlings. However, the variance in plant height did not reach a significant level between the control and PEG treatments (only slightly repressed by PEG treatment) (Figure 6F, G), which may have resulted from the mild stress intensity imposed by the low concentration of PEG6000. In *ct2* seedlings, the inhibition of growth by both the NaCl and PEG6000 treatments was more significant than that in WT (Figure 6F, G), indicating that *CT2* positively regulates salt and drought tolerance in maize. By contrast, the transcription of *CT2* was downregulated by the NaCl and PEG treatments (Figure 6H).

## Discussion

Heterotrimeric G protein complexes, which transduce signals from transmembrane receptors into cells, are key regulators of multiple fundamental cellular processes in plants [22]. Core G protein components and basic biochemistry are evolutionarily conserved among species. However, signaling mechanisms have been rewired (transmitting signals through atypical pathways and effector proteins) to satisfy the specific needs of plants[22, 39, 64]. G proteins have attracted the attention of agriculturalists mainly because of their functions in seed yield, biotic and abiotic stress responses, hormonal responses, nutrient usage, and symbiosis[39, 62]. However, studies on G protein signaling mechanisms have mainly focused on *Arabidopsis* and rice, and little is known regarding other crops such as maize.

In this study, we characterized a maize mutant, *ct2*, and cloned the obligatory target gene, *CT2*, through map-based cloning using the BC_1_F_1_ population. The mutation resulted from a 185 bp insertation at the carboxy-terminal of the *CT2* gene (Fig 2C). This insertion led to the replacement of 27 amino acids (aa) with a 19 aa sequence of CT2. Additionally, the transcriptional level of *ct2* was reduced in the *ct2* mutant (Fig 2B). The sequence alignment results showed that although there were three transversions in the promoters between WT and *ct2* (Fig S1), the promoter sequence of *ct2* in *ct2 mutant* is identical to that of B73. We speculate that the reduced expression of *ct2* was the result of the 185 bp insertation, and the mutant phenotype was from the inhibition of *CT2* expression or the mutation at the carboxy-terminal of *CT2* or both.

CT2 has been shown to physically interact with FEA2 to relay signals from ZmCLE7 to its downstream components to regulate the development of shoot meristems in maize[61]. The null mutation of *CT2* led to a short plant structure, shorter and wider leaves, wider meristems, strong fasciated ears, thicker tassel branches, and a higher density of spikelets [43]. Further studies showed that the mull mutation of *CT2* increased the total number of leaves, and *CT2* was involved in root system architecture and female inflorescence formation. More importantly, *CT2* suppressed the overproduction of female inflorescences[16]. By contrast, the expression of constitutively active *CT2* increased the kernel row number and reduced the leaf angle[15].

Plant height is a major breeding trait of maize. *ct2* mutant was a dwarf. Considering that the total number of leaves per plant was the same as that of the WT plants, the stem diameter and cell size in internode were equal to WT (Figure S4), in addition to the fact and that *CT2* modulates cell proliferation of shoots and roots in maize [16], we speculate that the reduced plant height in *ct2* resulted from the reduction in cell numbers, which may be resulted from retarded cell proliferation in the stalk internode of maize.

Our *ct2* mutant also exhibited defective development in male and female organs, including fasciated ears, compact and thicker rachis, and tassel branches (Figure 1, 3A-C, 3I-L), similar to the *CT2* mull mutant that Bommert *et al.* described[43]. Surveys and quantification of yield-related traits showed that ear length, kernel thickness, kernel length, and hundred-kernel weight significantly differed between *ct2* mutant and WT. Therefore, *CT2* is involved in yield-related traits. In addition, we found that the ear bracts were obviously shorter around *ct2* ears than in WT ears, and the tops of *ct2* ears (fasciated part of *ct2* ears) were poorly pollinated (Figure S5A). Notably, the tops of the ears in homozygous *ct2* BC_1_F_1_ plants were more fasciated than those in the parental inbred line *ct2* (Figure S5B). The ‘heterosis’ of the fasciation at the top of the mutant ears indicates that the mutant phenotype is affected by genetic background, or it merely reflects the result from the enlarged ears in hybrid. A functional study of *CT2* in different genetic backgrounds might answer the ‘heterosis’ phenomynon of the fasciation at the top of the mutant ears.

Although we have shown that the results are consistent with those in the literature, inconsistent results were also detected when compared with those of Bommert *et al.* [43], Urano *et al.* [16] and Wu *et al.* [15]. In these studies, the ct2 *null* mutant showed strongly fasciated ears and increased total leaves per plant, and constitutively active *ct2* increased the kernel row number. The ears of the *ct2* mutant characterized in this study were only slightly fasciated at the top, and mature ears were restored to normal, which may be attributed to poor pollination and the loss of the top part (Figure 3D, E). Total leaves per plant and kernel row number were uniformal to those of WT (Figure 3M, S4A). Notably, Bommert *et al.* and Urano *et al.* were *ct2* null mutant, and Wu *et al.* employed a transgenic line expressing constitutively active *ct2. ct2* mutant described in this study encoded a truncated ct2 protein with decreased transcriptional level of *ct2*. We speculate that truncated *ct2* retains part or complete function of CT2, and the inconsistent results were due to impaired protein function, reduced *ct2* expression, or both. The exonic expression of *ct2* and the mutants harboring weak *CT2* transcription levels will provide insights into inconsistent results with other studies.

G protein signaling exhibited conserved and rewired mechanisms in plants [33, 39, 59, 65–67]. Our phylogenetic analysis separates Gα into monocotyledonous and dicotyledonous groups, which indicates a divergency of Gα function in different crops (Figure 2D). Revealing the mechanism of Gα function in different plants is necessary. GO analysis showed that the functions of GPA1 (the *Arabidopsis* ortholog of CT2) interaction proteins included G protein-coupled receptor signaling, phytohormone (ABA and GA) signaling, blue light signaling, regulation of cell proliferation, and ROS metabolic processes (Table S5). The promoter sequence of *CT2* contains cis-regulating elements involved in signaling of phytohormones (ABA and MeJA), defense and stress responses, light responses, and meristem expression (Table S4). GO analysis of the DEGs between WT and *ct2* also detected genes involved in phytohormone (JA) regulation, signal transduction, stress, and defense responses (Figure 5). In *ct2,* the contents of JA-Ile was significantly depressed than in WT (Figure 6B). It is known that JA signaling pathways are involved in abiotic stress, including drought and salinity stresses, working in signal network with other phytohormone signaling pathways[68]. Salt and drought treatments in WT and *ct2* seedlings verified the involvement of *CT2* in stress tolerance and the feedback effect on *CT2* transcription (Figure 6F-G). Furthermore, both salt and drought treatments repressed the mRNA levels of *CT2*.

Based on the aforementioned results, *CT2* participates in maize biological processes (Figure 7). *CT2* is involved in various biological processes, including shoot meristem development, plant architecture, yield-related traits, JA signaling, and stress tolerance in maize. However, its underlying mechanisms remain revealing. For example, 1) the activity of G proteins relies on the formation and dissociation of the heterotrimeric complex between Gα and Gβγ, and we could not determine whether the mutant phenotypes of *ct2* resulted from the impaired function of *ct2* or the activation of Gβγ; 3) Does the altered salt and drought stress tolerance resulted from the decreased JA-Ile level in *ct2* or alternative pathway downstream of *CT2*? We are now creating transgenic lines of maize to overexpress and repress (or knock out) the expression of *CT2* to solve these questions. In conclusion, our results provided clues for the involvement of *CT2* in JA signaling and abiotic stresses, the resolving of underling mechanism will improve the G protein signaling pathway and helpful to screen beneficial alleles for maize breeding.

**Figure 7.**
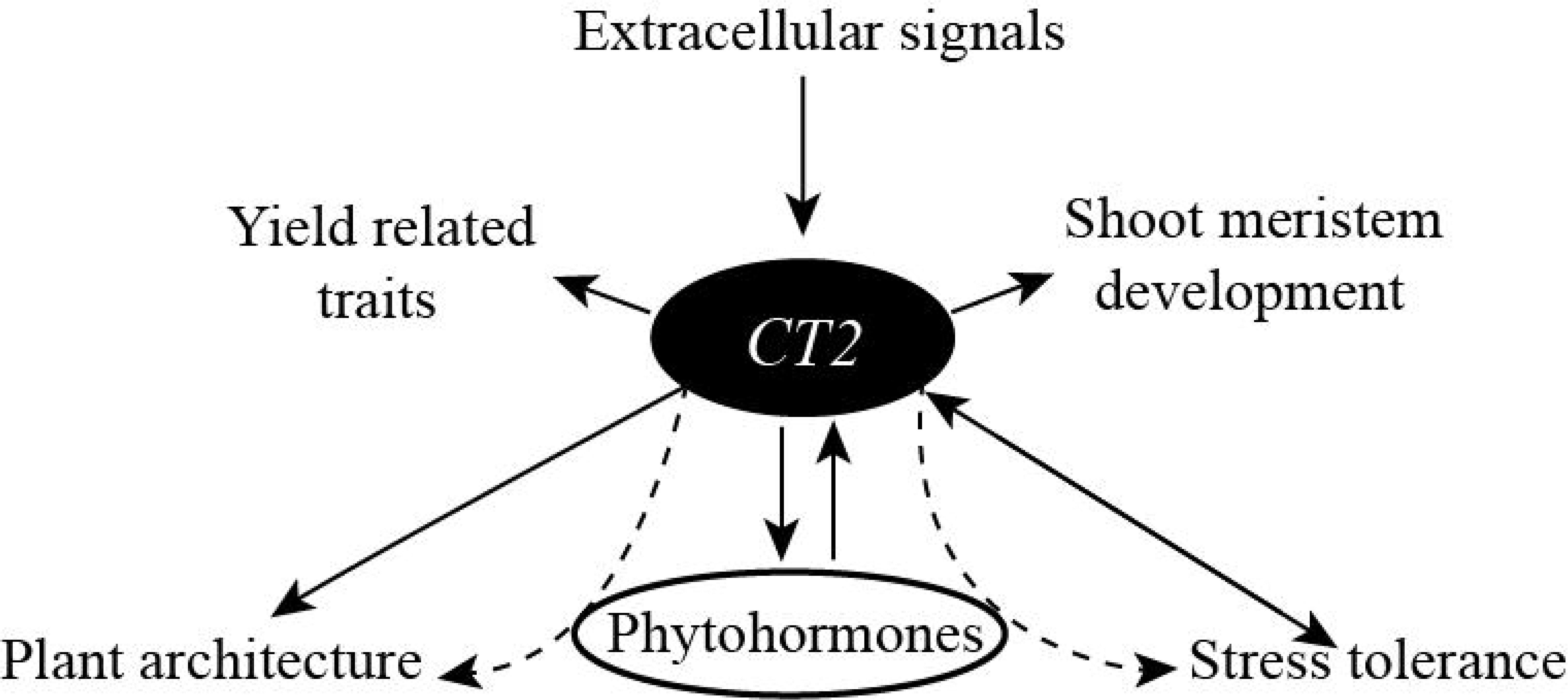
*CT2* participated biological processes in maize. Arrowheads shows the involvement of *CT2* in corresponding biological processes or the feedback effect to the expression of *CT2.* Solid and dotted lines indicate proven and putative processes.

## Supplemental information

### Supplemental Figures

Supplemental Figure S1. Sequence alignment of *CT2* promoters in WT and *ct2* mutant

Consensus sequences and mismatched nucleotide acids were highlighted in dark blue. and red rectangles, respectively. Arabic numerals at the right sides of the nucleotide acid sequences indicates the positions of the last nucleotide acid in each line relative to the adenine nucleotide in the translation initiation site ATG.

Supplemental Figure S2. Sequence alignment of *CT2* cDNAs in WT and *ct2* mutant Consensus sequences were highlighted in dark blue. The sequence of 185bp insertion was highlighted in blue with the inner terminator marked in red rectangles. Arabic numerals at the right sides of the nucleotide acid sequences indicates the positions of last nucleotide acid in each line relative to the adenine nucleotide in the translation initiation site ATG.

Supplemental Figure S3. Protein structure prediction of CT2 with InterPro Amnio acid sequence of CT2 was submitted to InterPro (http://www.ebi.ac.uk/interpro/) and analysed. The amnio acid sequence is indicated in red line. Arabic numerals on scale lines indicate amnio acid positions relative to the methionine at the translation initiation site, which is count as 1, of CT2. Black inverted triangle indicates the insertion site of the exogenous sequence in *ct2* mutant. Name of predicted protein domains were listed at the left side of the panel and the corresponding sequences were represented under the CT2 sequence.

Supplemental Figure S4. Quantification of stem diameter and numbers of leaves per plant in WT and ct2 *mutant*

Stem diameter and numbers of leaves per plant were investigated at the mature stage in WT and ct2 *mutant.* At least ten plants were investigated and averaged for each trait.

**A**, Quantification of stem diameter.

**B**, Quantification of numbers of leaves per plant.

Error bars represent stand deviation errors. ns, not significant. **, P< 0.01.

Supplemental Figure S5. Ear related phenotype in WT and *ct2* mutant

**A**, Ears of 25 days after pollination with ear bract retained (upper panel) or stripped (lower panel) of WT and *ct2* mutant.

**B**, WT and *ct2* mutant type ears before pollination (upper panel) or 25 days after pollination (lower panel).

### Supplemental tables

Table S1. BSA mapping result of the candidate gene responsible for the mutant phenotype of *ct2*.

Table S2. Primers used in this study

This table lists the primers used in this study.

Table S3. RNA-seq analysis for WT and *ct2 mutant*

This table lists the differentially expressed genes attained from the RNA-seq data analysis of leaves in WT and *ct2* mutant.

Table S4. *Cis*-acting regulatory elements on promoter of *CT2*

This table lists cis-acting regulatory elements predicted by Plant CARE website (http://bioinformatics.psb.ugent.be/webtools/plantcare/html/) on promoter (−2320 - 0bp before the translation initiation site ATG) of *CT2*.

Table S5. Gene ontology analysis of GPA1 interacting proteins

Result of gene ontology analysis of GPA1 interacting proteins were obtained from BioGRID ORCS (https://thebiogrid.org/2522/summary/arabidopsis-thaliana/gp-alpha-1.html) and listed in the table.

## Author contribution statement

WJ, JY, JT, and XZ designed the study. YS and WC participated in the experiments. QZ and ZG analyzed the BSA-seq and RNA-seq data. LY and XY mainly participated in the phenotyping and map-base cloning of the candidate genes, respectively. YL performed an experiment to determine the effect of phytohormones on the expression of *CT2*. KZ and LZ were responsible for the stress treatment experiments. The manuscript was written mainly by YS, with the assistance of WJ, JY, JT, and XZ. All the authors have read and approved the final version of the manuscript.

## Funding

This work was supported by the Natural Science Foundation of Henan Province (No. 232102111097), the Science and Technology Innovation Fund of Henan Agricultural University (No. KJCX2021A01).

## Data availability

Datasets generated during the current study are available from the corresponding author on reasonable request.

## Conflict of interest

The authors declare that there is no conflict of interest.

## Notes

### Competing Interest Statement

The authors have declared no competing interest.

