## Supplementary figures and images for "Map based cloning of *CT2* and the pilot functional exploration in abiotic stress"

### Supplemental Data 1

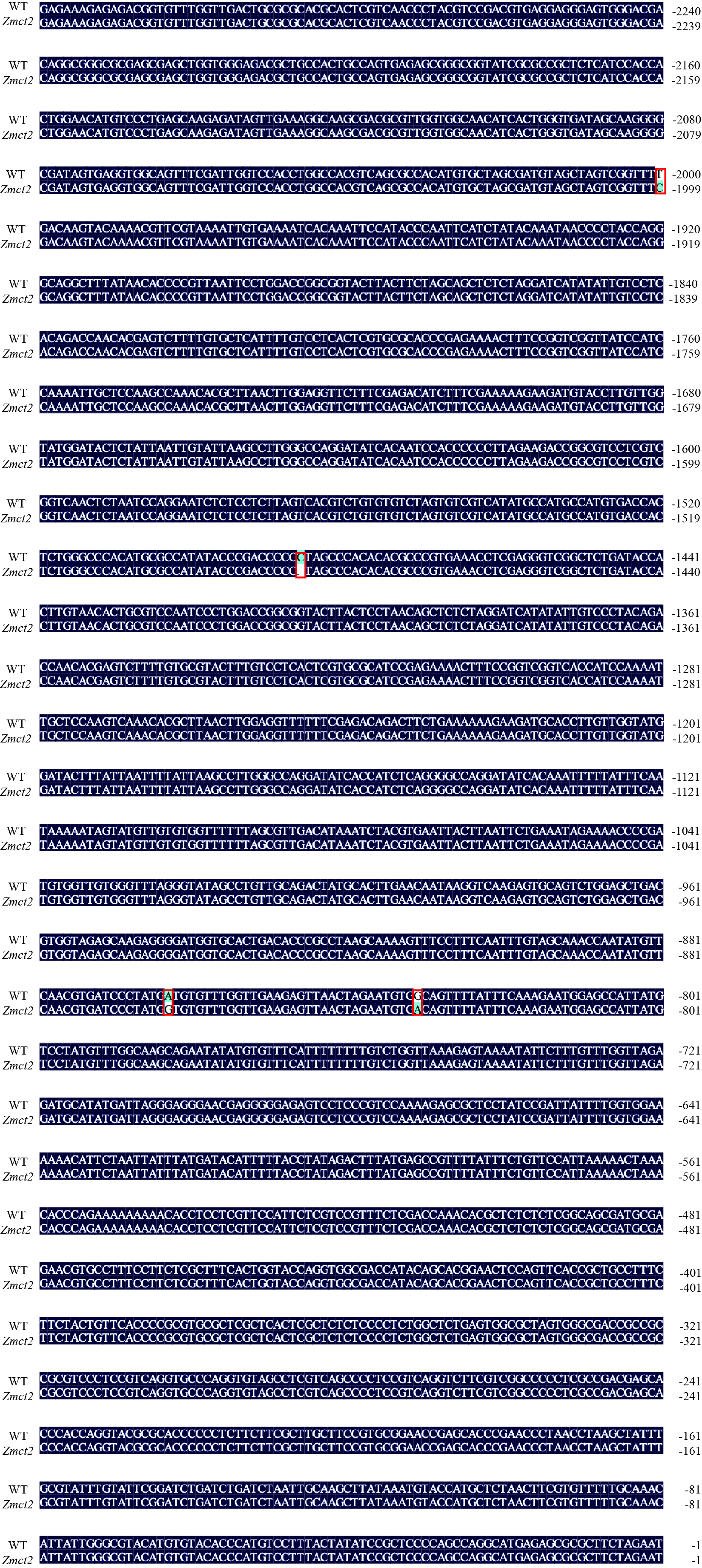

### Supplemental Data 2

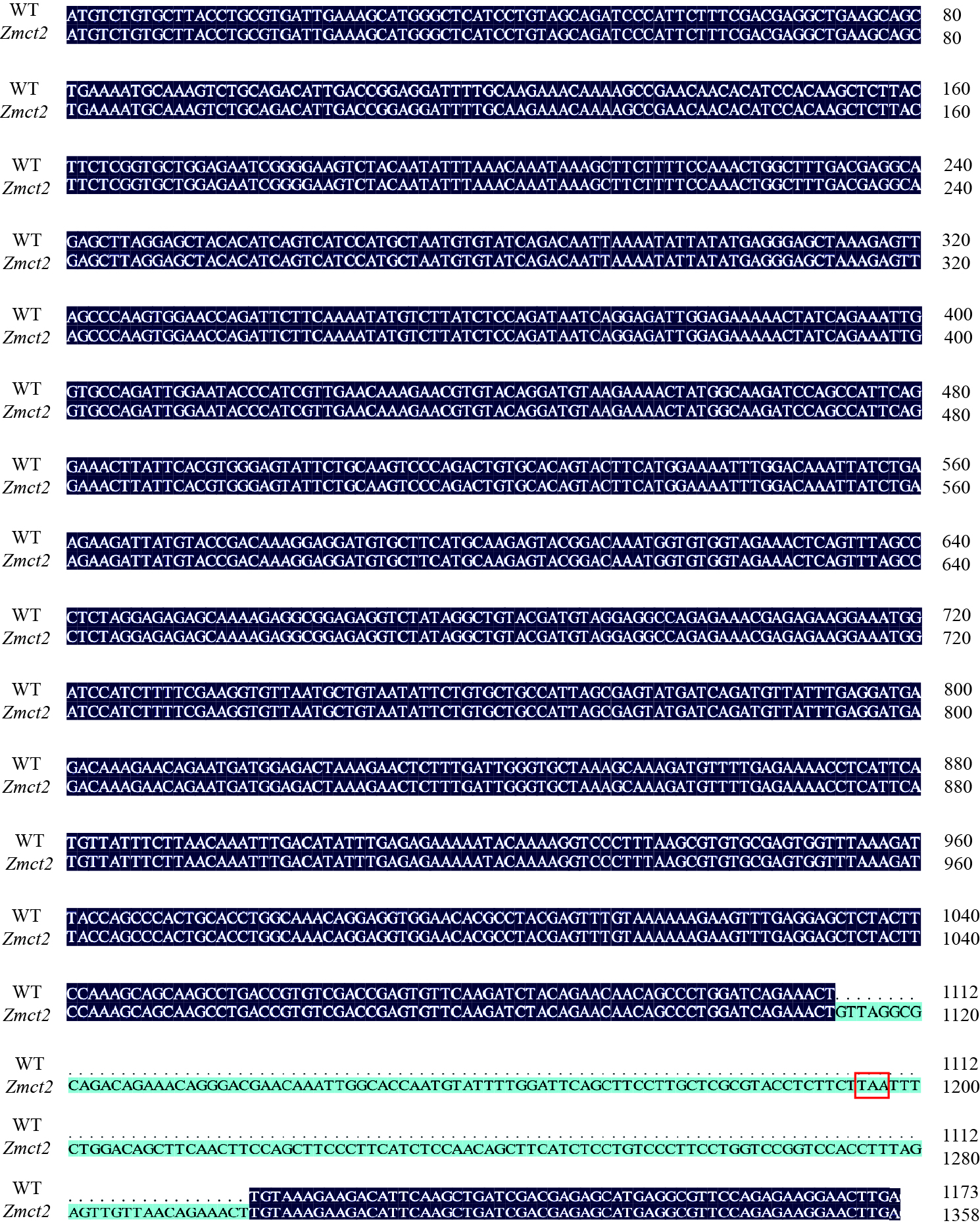

### Supplemental Data 3

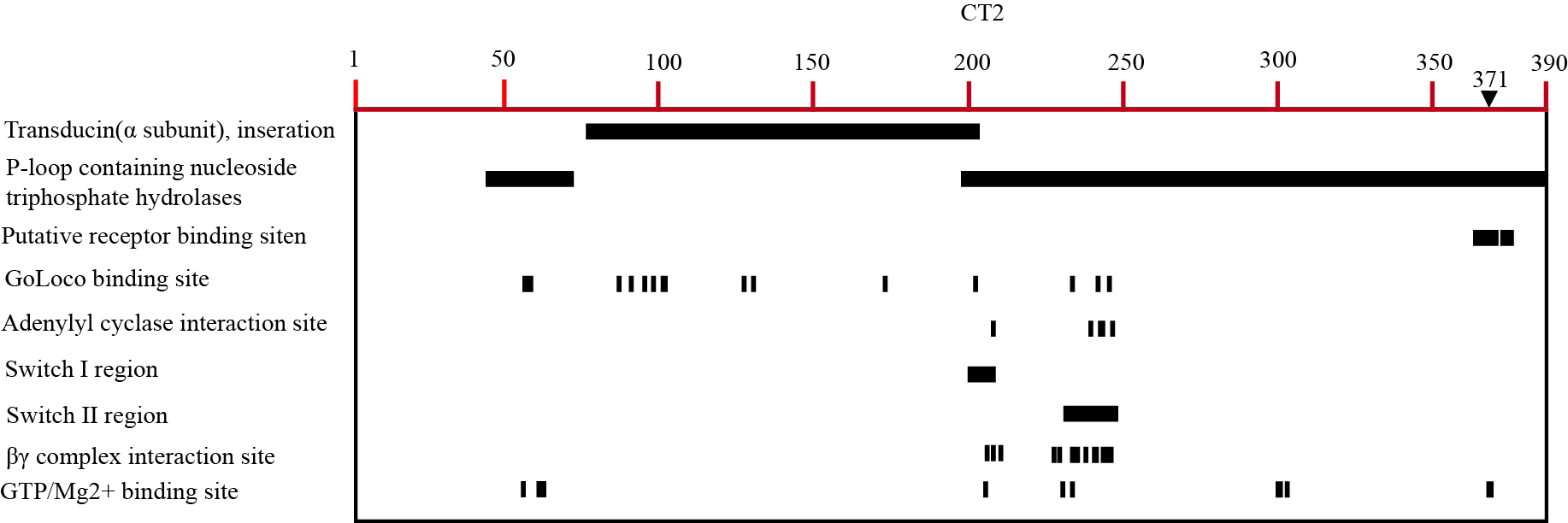

### Supplemental Data 4

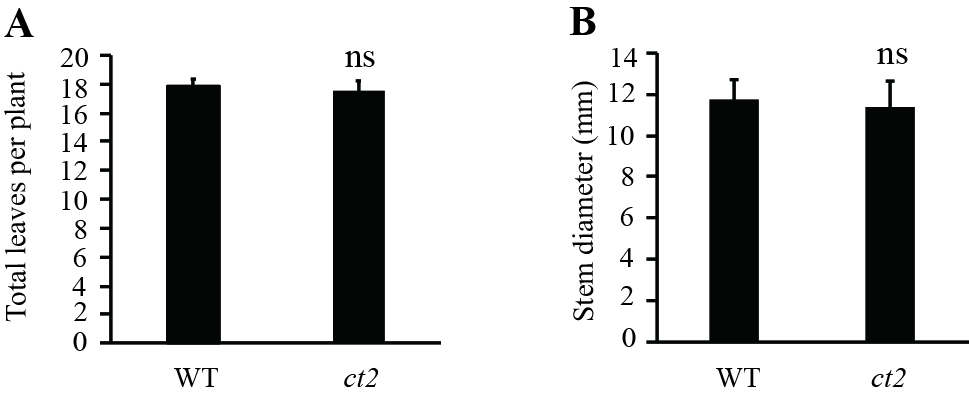
